# Telomeric Repeat-Containing lncRNA TERRA Targets Non-Telomeric DNA in Trans via R-Loops

**DOI:** 10.1101/2025.06.29.662205

**Authors:** Saifeldin N. Shehata, Akram Mendez, Tanmoy Mondal, Roshan Vaid

## Abstract

Telomeres are transcribed into telomeric repeat-containing RNA (TERRA), a long non-coding RNA known to play key roles in telomere maintenance and DNA repair. While TERRA’s formation of R-loops at telomeric regions is well documented, its interaction with non-telomeric DNA remains poorly understood. In this study, we reanalyzed publicly available datasets using bioinformatic approaches and found evidence that TERRA binds non-telomeric DNA regions in trans through R-loop formation. Notably, the presence of tandem telomeric repeat motifs within non-telomeric regions enhanced TERRA binding, suggesting sequence-dependent R-loop formation. Furthermore, analysis of previously identified TERRA-regulated genes revealed that such distal TERRA-DNA interactions may influence telomere elongation by modulating the expression of telomere maintenance factors. These findings provide new insights into the broader regulatory functions of TERRA beyond telomeres and suggest a potential mechanism for its role in genome-wide gene regulation via R-loops.

## INTRODUCTION

TERRA (telomeric repeat-containing RNA), a long non-coding (lnc) RNA originating from sub-telomeric regions extending towards telomere ends, was first reported 25 years ago in *Trypanosoma brucei*^*1*^. In humans, TERRA contains UUAGGG telomeric repeats and was first reported in 2008. TERRA is a heterogeneous population of lncRNAs, varying widely in length (100 bases to >9 kb)^2^, and may or may not possess polyA tails in the mature RNA transcripts^3^. Despite over two decades of study, many aspects of TERRA remain to be understood.

Telomere maintenance primarily depends on the activity of the telomerase enzyme encoded by the TERT gene. However, a telomerase-independent alternative mechanism known as Alternative Lengthening of Telomeres (ALT) also exists. Although the exact mechanisms driving cells towards ALT are not fully understood, inactivating mutations in the alpha thalassemia/mental retardation syndrome X-linked (ATRX) gene are thought to be a key factor. We and others have shown that TERRA is highly expressed in various cancers positive for ALT^4-6^. This aligns with the notion that ATRX antagonizes TERRA^7^. Studies in yeast where TERRA was upregulated revealed cells with reduced senescence and longer telomeres^8,9^. TERRA forms R-loops (RNA-DNA hybrid structures) in cis at the telomere ends where it is transcribed, playing critical roles in telomere maintenance and elongation^4,10-13^.

While telomere maintenance by TERRA suggests its cis-acting function, it has been proposed that TERRA also functions in trans. Using TERRA CHIRT sequencing (Chromatin Isolation by RNA Purification, ChIRP, Capture Hybridization Analysis of RNA Targets, CHART), TERRA binding was mapped across the entire mouse genome. Interestingly, TERRA binding to non-telomeric regions of the genome was also identified in a significant proportion^7^. This suggests either a role for TERRA outside of telomere maintenance or that TERRA exerts telomere maintenance by binding non-telomeric regions of the genome.

Using a TERRA reporter system, Feretzaki et al. demonstrated that trans-acting TERRA is targeted to telomere ends by forming R-loop structures with the help of proteins such as RAD51 and BRCA2^13^. Importantly, how TERRA interacts with its non-telomeric genomic targets in trans remains largely unknown. To the best of our knowledge, there is only one report to date in *Arabidopsis thaliana* demonstrating a lncRNA, APOLO, acting in trans via R-loop formation and regulating expression of distal genes^14^. In this study, by bioinformatically reanalyzing genome-wide R-loop (DNA-RNA immunoprecipitation followed by sequencing, DRIP-seq) and TERRA binding (TERRA CHIRT-seq) data in mouse embryonic stem cells, we explored if trans TERRA binding is driven by R-loops. This could be a prevalent mechanism employed by other trans-acting lncRNAs to modulate the expression of their target genes.

## MATERIAL AND METHODS

### Raw data accession numbers and data visualization

TERRA CHIRT-Seq (anti-sense: SRR2062968, sense control: SRR2062969), R-loop DRIP-Seq (SRR2075686), and ATRX ChIP-Seq (SRR057567) sequencing files were obtained from the Short-Read Archive (SRA) database^15^ using the Gene Expression Omnibus (GEO) accessions for TERRA (GSE79180), R-loop (GSE70189), and ATRX (GSE22162) using the Entrez Direct E-utilities^16^.

All bar-plots, boxplots, karyo-plots and Venn diagrams were generated using ggplot2 v3.3.6 and KaryoplotR v1.18.0 packages in R^17,18^. All metagene plots were generated using DeepTools v3.5.1 (computeMatrix, plotProfile, and plotHeatmap)^19^. All peaks were visualized using Gviz v1.36.2 (https://bioconductor.org/packages/release/bioc/vignettes/Gviz/inst/doc/Gviz.html, https://github.com/ivanek/Gviz) in R. Motif analysis was performed using MEME v5.4.1^20,21^, and pie-charts were generated using both ggplot2 and Microsoft Excel.

### Identifying TERRA and R-loop peaks

Sequencing reads in FastQ format were retrieved from the SRA database using Sra-Toolkit v2.11.0 (fastq-dump) developed by the SRA Toolkit Development Team at^22^. After quality control using FastQC v0.11.9^23^, sequences with a Per Base Sequence Quality of less than 28 were trimmed using Trimmomatic v0.39^24^. The resulting reads were aligned to the mouse mm10 genome using BWA-MEM v0.7.17^25^, and peaks were identified using MACS2 (callpeak) v2.2.7.1^26,27^.

### Identifying overlapping peaks

Narrow Peak files from MACS2 peak calling for both TERRA and R-loop were used as input to BEDTools v2.30.0 (intersectBed)^28^ to obtain peaks that contained a minimum of one base pair overlap/intersection.

### Annotating peaks and counting telomeric repeats per peak

The overlapping TERRA peak file (generated with intersectBed) was annotated using HOMER v4.11 (annotatePeaks.pl) with specific parameters (-nmotifs and -mbed)^29,30^ to identify the location and number of telomeric repeats in each peak. The resulting annotation file was used to count and plot the number of repeats per peak in R.

### Extracting peaks with tandem repeats

Using the HOMER-generated annotation file as input, a custom Bash/Command Line script was used to identify and extract only the annotated peaks containing a minimum of four tandem repeats.

### Coverage profiles and heatmaps

Read coverage was computed using computeMatrix, plotProfile and plotHeatmap from DeepTools (REF: https://deeptools.readthedocs.io/en/latest/). Alignment bigwig files were used as input score files and intersecting peak BED files as the input regions files. Regions files included R-loop peaks that intersected with TERRA peaks split by presence or absence of tandem telomeric repeats, as well as by genome annotation (intron or intergenic). Peaks that lay within the ENCODE blacklisted regions (https://github.com/Boyle-Lab/Blacklist/raw/master/lists/Blacklist_v1/mm10-blacklist.bed.gz) or that were within 500kb from telomeres were excluded from the resulting profiles and heatmaps.

## RESULTS

### A small fraction of TERRA and R-loop peaks overlapped

To determine whether TERRA RNA binds to non-telomeric DNA sites in trans via R-loops, we obtained published datasets containing sequencing reads after TERRA^7^ or R-loop^31,32^ immunoprecipitation from mouse embryonic stem cells. After processing and aligning the reads to the mouse (mm10) genome, we identified peaks to determine the genomic coordinates where TERRA and R-loops are most likely located. To identify the maximum number of potential loci, we chose not to filter the peaks by fold enrichment, resulting in approximately 28,000 TERRA peaks and 29,000 R-loop peaks (Fig. 1A). Notably, filtering for peaks with a fold enrichment of at least 10 over control resulted in 4,000 TERRA peaks, confirming results by Chu et al ^7^.

**Figure 1.**
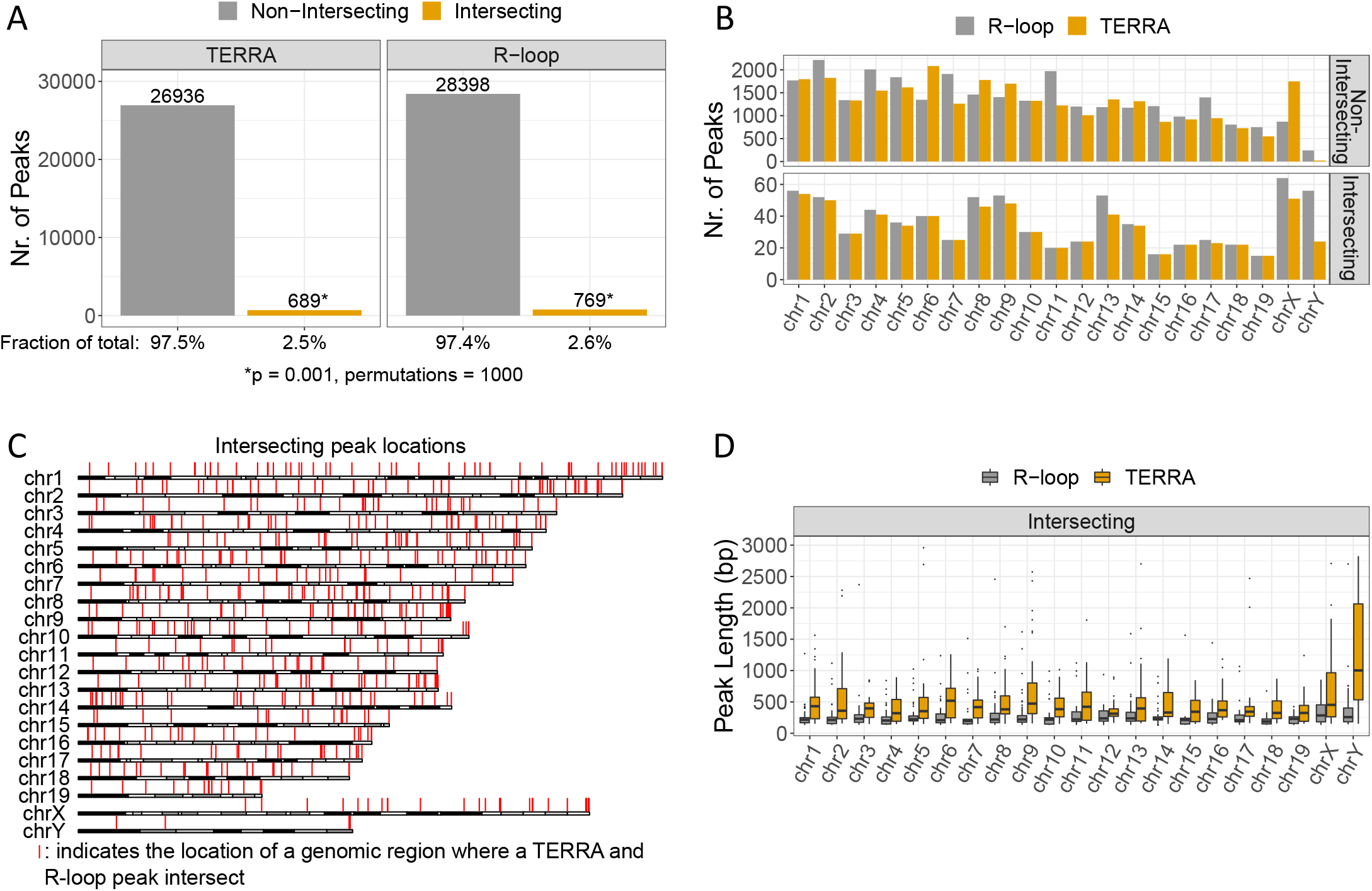
A small fraction of TERRA and R-loop peaks overlap. **A)** Total number of TERRA and R-loop peaks, including the fractions of non-intersecting and intersecting peaks. A hypermutation test was performed to assess statistical significance. **B)** Number of peaks per chromosome by peak type (TERRA vs. R-loop) and group (non-intersecting vs. intersecting). **C)** Genomic positions of intersecting peaks displayed across each chromosome. **D)** Distribution of peak lengths per chromosome for each peak type.

If TERRA binds its targets via R-loops, their locations on the genome should coincide, leading to overlapping/intersecting peaks in genomic regions. To maximize the number of intersecting peaks, we filtered for TERRA and R-loop peaks that contained an intersection of at least one base pair. Only a small fraction (less than 3%) of peaks intersected (Fig. 1A), suggesting alternative mechanisms for TERRA binding, which may reflect a diversity in TERRA binding modes and physiological roles. To confirm whether the observed intersection between TERRA and R-loop peaks was significant, we performed a hypermutation test with randomized peak regions and found the intersections to be highly significant (n = 1000 permutations, p < 0.001, Fig. 1A and S1A).

The significance of intersecting peaks prompted further investigation. We assessed the distribution of intersecting peaks across all chromosomes and found that they were present in comparable numbers across all chromosomes, precluding associations with any specific chromosome (Fig. 1B). Since TERRA originates at telomeres, we examined whether the majority of intersecting peaks were enriched at telomeric or sub-telomeric regions. Analyzing the intersecting peak locations on individual chromosomes revealed that although there was some enrichment at the ends of select chromosomes (e.g., 9, 10, 11, and 13), intersecting peaks were distributed across each chromosome (Fig. 1C).

R-loops are typically associated with co-transcriptional activity^33^, which, depending on gene length, may lead to broad R-loop peaks spanning long stretches of DNA along transcribed regions. We assessed the average R-loop peak length by comparing TERRA and R-loop peak lengths per chromosome, and found that R-loop peaks were on average very narrow (∼150-350bp) (Fig. 1D), which may support a distal/trans binding event of TERRA rather than a co-transcriptional one. Interestingly, intersecting TERRA peaks on all chromosomes were on average larger than intersecting R-loop peaks (Fig. 1D). Notably, many R-loop peaks lay completely within TERRA peak regions, and some TERRA peaks spanned regions containing more than one R-loop peak (data not shown).

### Telomeric repeats are highly enriched in intersecting peaks

A defining feature of TERRA is the presence of long stretches of short tandem telomeric repeat sequences (5’-UUAGGG-3’). However, telomeric repeats are also found at non-telomeric sites across the genome^7^. The presence of telomeric repeats in both TERRA and genomic regions should, in principle, facilitate R-loop formation due to sequence complementarity. To investigate this, we counted the total number of single telomeric repeats in all TERRA-bound peaks and calculated their distribution across R-loop-intersecting vs. non-intersecting peaks. Surprisingly, 44% of all repeats were enriched in the intersecting peaks, which only represented 2.5% of all TERRA binding regions, while the remaining non-intersecting peaks (97.5% of all TERRA peaks) harbored 56% of telomeric repeats (Fig. 2A). This suggested that TERRA RNA targeting in trans is achieved via telomeric repeat-dependent R-loop formation.

**Figure 2.**
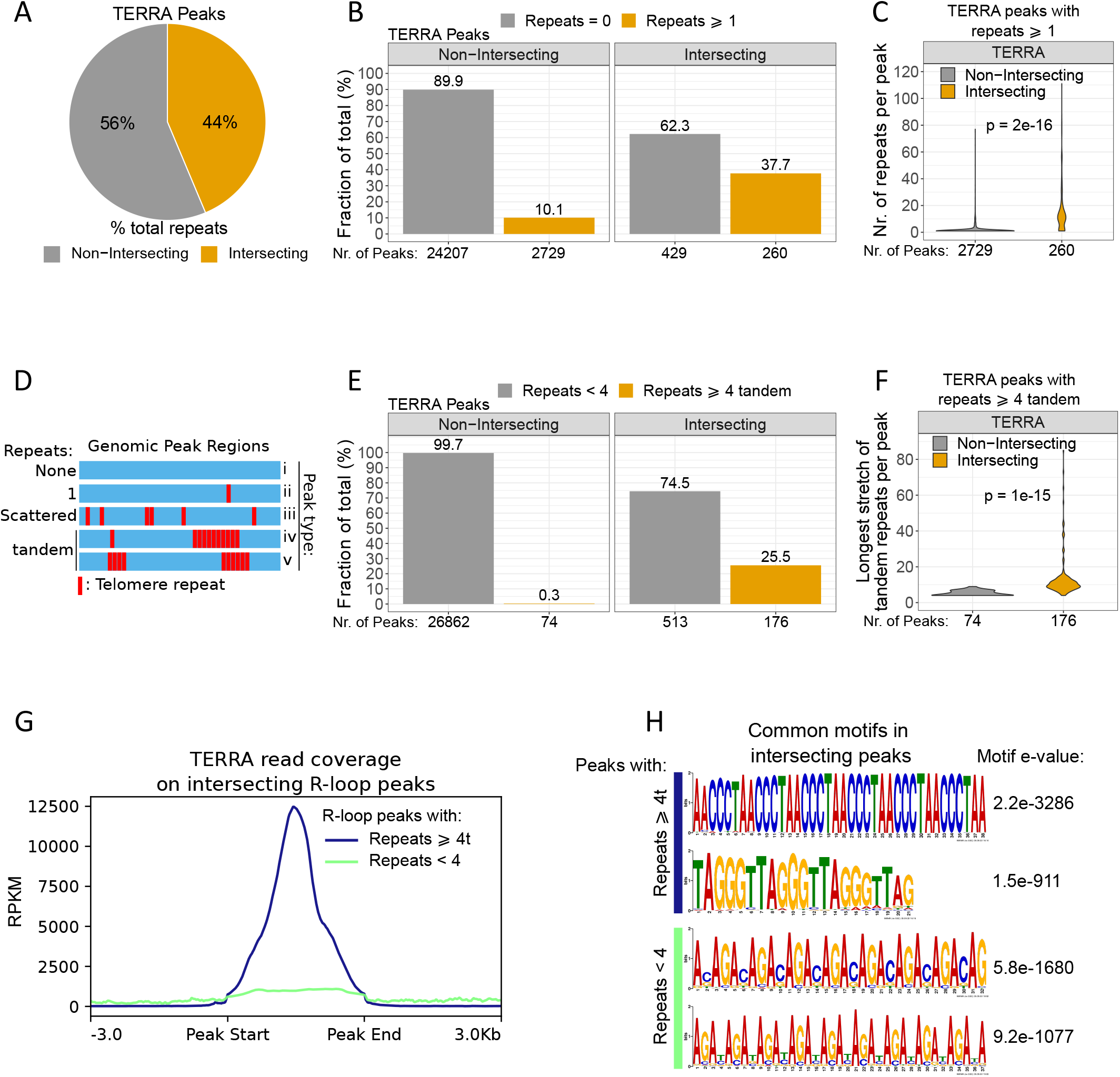
Telomeric repeats are highly enriched in overlapping peaks. **A)** Fraction of intersecting and non-intersecting TERRA peaks that harbour at least one telomeric repeat. **B)** Fraction of intersecting and non-intersecting peak groups that either contain or lack at least one telomeric repeat. **C)** Distribution of telomeric repeat counts per TERRA peak, separated into intersecting and non-intersecting groups. Only peaks containing more than one telomeric repeat were included. **D)** Categorization of peaks into five groups based on the number and distribution of telomeric repeats. **E)** Fraction of intersecting and non-intersecting TERRA peaks containing fewer than four versus at least four tandem telomeric repeats. **F)** Distribution of the longest stretch of telomeric repeats per TERRA peak, separated into intersecting and non-intersecting groups. Only peaks with at least four tandem telomeric repeats were included. **G)** Coverage profiles of TERRA sequencing reads over genomic regions intersecting with R-loops, separated into peaks with fewer than four versus at least four tandem telomeric repeats. **H)** Most common sequence motifs in TERRA/R-loop intersecting peaks containing fewer than four versus at least four tandem telomeric repeats.

Next, we filtered TERRA peaks that contained at least one telomeric repeat. We found that while ∼38% of intersecting peaks (260 out of 689 peaks) contained at least one repeat, only ∼10% of non-intersecting TERRA peaks (2729 out of 26936 peaks) contained at least one repeat (Fig. 2B). We then investigated the frequency of these telomeric repeats per TERRA peak. Interestingly, the majority of the intersecting TERRA peaks (82%, 212 of 260 peaks) had more than one telomeric repeat, with a median of 11 repeats per peak. On the other hand, close to 81% (2215 of 2729) of non-intersecting peaks had only one telomeric repeat (Fig. 2C), with a median of one repeat per peak, suggesting that TERRA peaks that intersect with R-loops have a distinct enrichment of consecutive/tandem telomeric repeats.

To explore this further, we assessed the repeats within all TERRA peaks and divided them into five peak types (Fig. 2D). Considering the in vivo requirements for R-loop initiation and stabilization, we hypothesized that a minimum of four tandem telomeric repeats (i.e. TTAGGG x 4 or CCCTAA x 4) would be needed for the formation of stable R-loops (Fig. 2D, peak types iv & v). We therefore excluded all TERRA peaks lacking a stretch of at least four tandem telomeric repeats. We found that while a quarter (∼25%) of intersecting peaks passed this filter, the vast majority (99.7%) of non-intersecting peaks lacked tandem repeats (Fig. 2E), further confirming the high enrichment of long stretches of telomeric repeats only in TERRA peaks that intersected with R-loops. To further validate this, we counted the longest stretch of tandem repeats in each TERRA peak, while excluding peaks with less than four tandem repeats, and found that only intersecting TERRA peaks contained long stretches of tandem repeats (Fig. 2F). This confirms that although many non-intersecting TERRA peaks contained a high number of telomeric repeats (Fig. 2C), the vast majority of these repeats are scattered (Fig. 2D, peak type iii) and are likely unable to form R-loops in vivo. This data suggests that the presence of tandem telomeric repeats in genomic DNA can serve as a predictor of TERRA-dependent R-loop formation.

We noted that ∼75% of R-loop-intersecting TERRA peaks (513 out of 689) did not contain tandem telomeric repeats (Fig. 2E), which prompted further investigation. We asked if the presence of telomeric repeats in the intersecting TERRA peaks correlated with TERRA enrichment, and found that TERRA enrichment was 10-fold higher in peaks with four or more tandem repeats (Fig. 2G). This indicated a preference for TERRA binding to telomeric repeat-enriched genomic regions. Furthermore, to assess the effectiveness of our four-tandem-repeat filter and to understand why many TERRA peaks lacking tandem repeats still intersected with R-loop peaks, we performed a motif analysis on the peak regions. This analysis confirmed that telomeric repeats (TTAGGG or CCCTAA) were only found in those peaks that passed our filter, while peaks that lacked tandem telomeric repeats were enriched in AG-rich motifs (Fig. 2H). Collectively, these results implicate the presence of stretches of tandem telomeric repeats as predictors of TERRA-dependent R-loop formation.

### ATRX coverage is analogous with TERRA

The regulation of gene transcription is a complex process involving many factors, and it is conventionally thought to be regulated primarily by transcription factor binding to promoter sequences^34^. Another major regulator of gene transcription is enhancer sequences, which can be up to tens of megabases away from the gene promoter yet still regulate transcription^35^. We observed that the majority (over 90%) of R-loop-intersecting TERRA peaks were located either within introns (∼43%) or in intergenic regions (∼51%) (Fig. 3A). When assessing the relative enrichment of TERRA sequencing reads on intergenic and intronic R-loop regions, we found that peak regions with tandem repeats had significantly higher coverage than peak regions lacking them, regardless of whether the peaks were within introns or intergenic regions (Fig. 3B, left panel).

**Figure 3.**
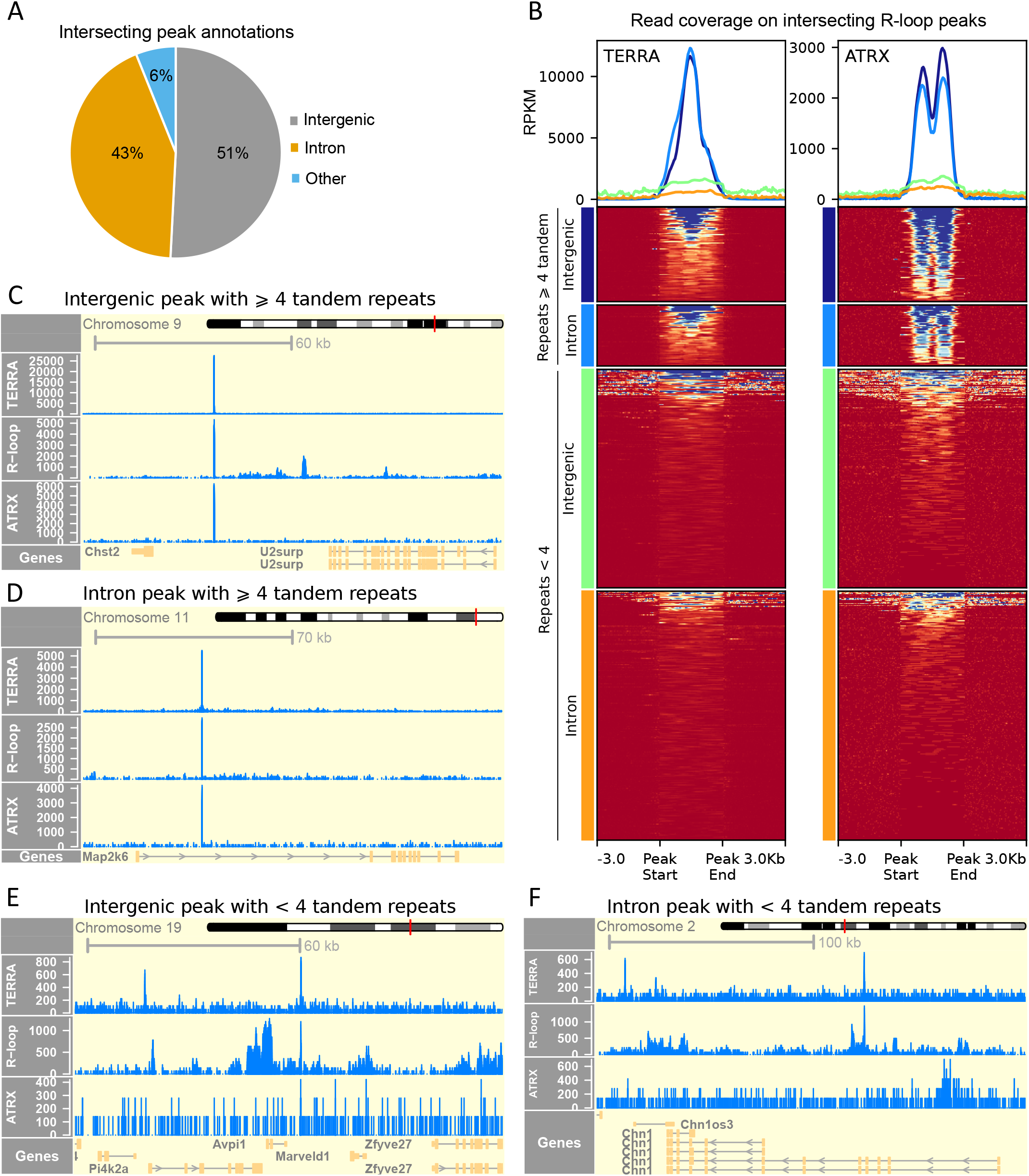
ATRX overlaps with TERRA-R-loop peaks, but not within peaks lacking tandem telomeric repeats. **A)** Genomic distribution of TERRA/R-loop intersecting peaks, categorized into intronic, intergenic, or other regions (promoter–TSS, TTS, 3′UTR, exons). **B)** Metagene profiles and heatmaps of TERRA and ATRX coverage over intersecting peaks located in intronic or intergenic regions. Peaks were further divided into groups containing fewer than four versus at least four tandem telomeric repeats. **C–D)** Genome browser views of TERRA, R-loop, and ATRX coverage at intergenic (C) and intronic (D) peaks containing at least four tandem telomeric repeats. **E–F)** Genome browser views of TERRA, R-loop, and ATRX coverage at intergenic (E) and intronic (F) peaks containing fewer than four tandem telomeric repeats.

Furthermore, it was recently shown that ATRX, a chromatin remodeler, interacted with TERRA at many genomic loci^7^, where the authors demonstrated that ATRX competes with TERRA for DNA binding to regulate telomere maintenance and protect telomeres^7^. If ATRX indeed antagonizes TERRA, then it is expected to be present where TERRA binds and would have a matching coverage profile to TERRA on intersecting R-loop peaks. We utilized a publicly available ATRX ChIP-seq dataset^36^ and found a high relative enrichment of ATRX ChIP-seq reads on the same genomic regions as telomeric repeat-enriched TERRA peaks. Indeed, we observed an analogous ATRX profile to that of TERRA (Fig. 3B, right panel), with a high presence of ATRX only in repeat-enriched peaks. The associated heatmap shows each peak as a horizontal line, with a blue color indicating higher enrichment in that peak region. The heatmap clearly shows that the presence of telomeric repeats, rather than genome annotation, is the main factor associated with ATRX presence at those sites.

This strongly indicated that the R-loops detected at these genomic regions have not originated co-transcriptionally but were rather due to the trans binding of TERRA originally transcribed at the telomeres and migrating to non-telomeric genomic loci to form R-loops.

To further validate this, we visualized individual intersecting TERRA peaks that lay within telomeric repeat-enriched genomic regions and observed stronger signals for TERRA, R-loops, and ATRX peaks in both intronic and intergenic regions (Fig. 3C and D) compared to those that were not repeat-enriched (Fig. 3E and F). We also observed broad R-loop peaks spanning several exons within genes, for instance in the Death-associated protein kinase 1 (Dapk1) and BRCA1-associated RING domain protein 1 (Bard1) genes (Fig. S3B and C). Interestingly, in the peaks within Dapk1 and Bard1, the R-loops associated with TERRA, though surrounded by broader R-loop regions, were more highly enriched and spanned a much narrower region. This distinction from the surrounding R-loops further indicated that these specific R-loops are a result of the trans binding of TERRA rather than co-transcriptional R-loops, supporting our hypothesis that TERRA binds non-telomeric DNA via R-loops.

### A small subset of genes harboring TERRA-R-loop intersecting peaks are differentially expressed upon TERRA knock-down

It was previously reported that several genes were differentially expressed upon TERRA knockdown^7^ Although the downstream effects of TERRA binding to DNA/protein involve adapter functions^32^ and may not be limited to direct modulation of gene transcription, we wanted to explore whether any genes associated with our R-loop-intersecting TERRA peaks were previously reported to be differentially expressed upon TERRA knockdown. Indeed, when we extracted genes associated with or in close proximity to telomeric repeat-enriched intersecting peaks, we observed a small subset of genes overlapped with those previously reported as differentially expressed in a TERRA-dependent manner (Fig. 4A).

**Figure 4.**
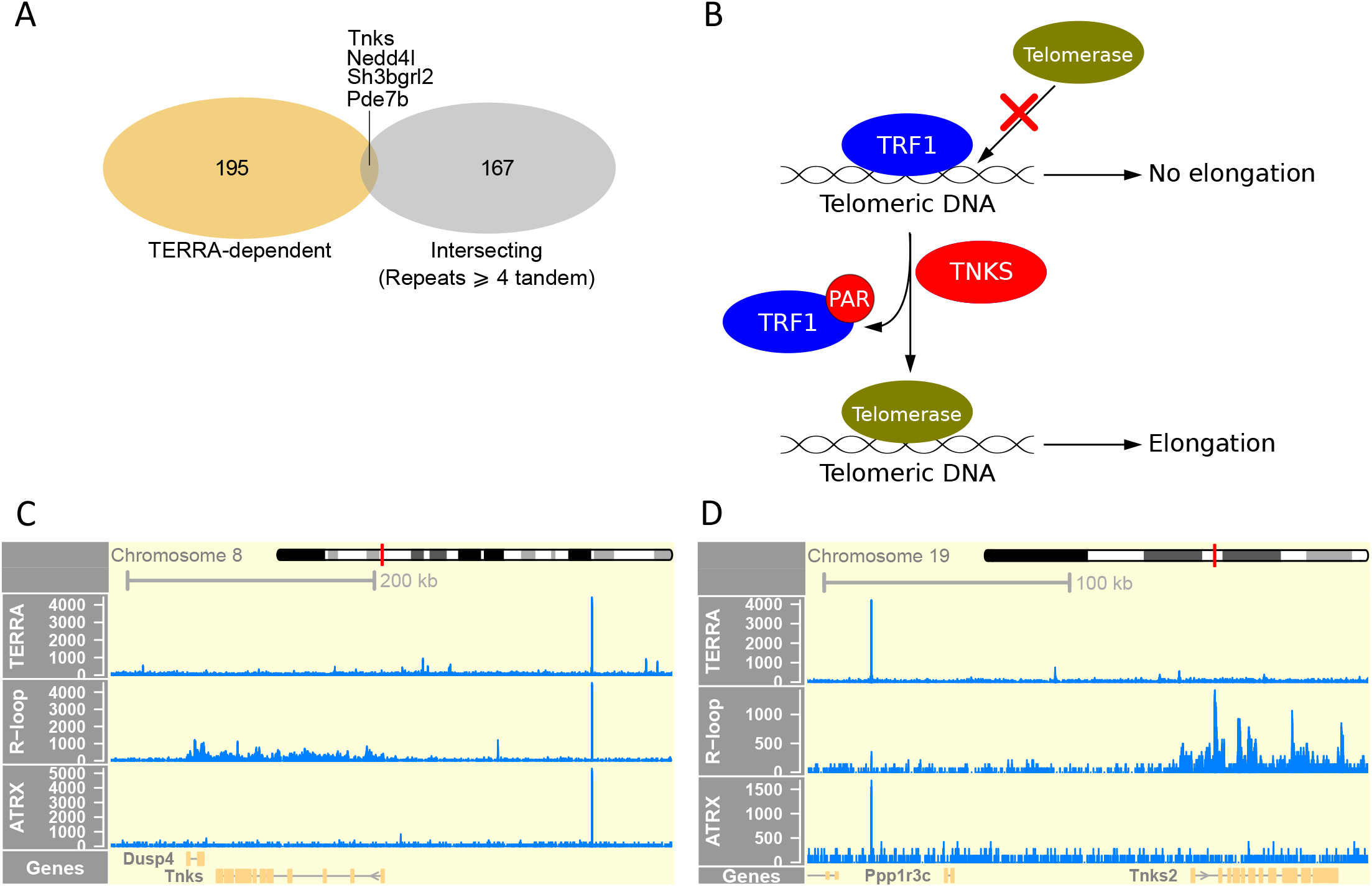
A portion of genes downregulated upon TERRA knockdown overlap with R-loops and harbour tandem telomeric repeats. **A)** Overlap between genes downregulated upon TERRA knockdown and genes containing TERRA/R-loop intersecting peaks within introns or in close proximity to the gene. **B)** Overview of the opposing effects of TRF1 (TERF1) and TNKS1 on telomere elongation. **C–D)** Genome browser views of TERRA, R-loop, and ATRX coverage at selected genes from the overlap in panel A.

Interestingly, one of these genes was Tankyrase 1 (TNKS1), a Poly-ADP-Ribosylation polymerase (PARP) known to regulate telomere elongation by PARsylation of Telomeric Repeat Binding Factor 1 (TRF1)^37-40^. PARsylation of TRF1 results in its release from telomeric DNA and subsequent degradation via ubiquitylation, allowing telomerase to elongate telomeres (Fig. 4B)^41,42^. It would be interesting to explore whether TERRA-dependent regulation of TNKS1 expression in turn affects telomere length and under which physiological conditions this may occur.

Moreover, both TNKS1 and TNKS2 are involved in telomere regulation, and we detected TERRA-R-loop-ATRX peaks within 130-170 kb of their promoter regions (Fig. 4C and D), which might implicate TERRA in the regulation of their expression.

Taken together, our results demonstrate a possible mechanism by which TERRA RNA acts in trans by forming R-loop structures at genomic regions that have tandem telomeric repeats, and that TERRA RNA knockdown may influence the expression of genes that harbor these R-loops in their vicinity. These findings further support the idea that such R-loop–mediated regulation could represent a more general mechanism employed by other trans-acting lncRNAs to modulate the expression of their target genes.

## DISCUSSION

In this study, we presented evidence that the presence of TERRA (Telomeric Repeat-containing RNA) at non-telomeric genomic sites rich in tandem telomeric repeats is highly correlated with R-loops. This suggests that TERRA can bind DNA in trans via R-loops, far from the telomere. We further show that this binding may partially explain previously reported changes in the expression of certain genes upon TERRA knockdown^7^. This implies that TERRA may function as a feedback mechanism, where it is transcribed at telomeres and then binds non-telomeric DNA in trans via R-loops to affect the expression of genes involved in telomere maintenance.

It would be interesting to identify the proteins involved in assisting with strand invasion to form these R-loops at non-telomeric genomic sites, and whether they are different from those that assist TERRA-R-loop formation at telomeres. A recent report showed how ATRX binding to RNAs can recruit PRC2 in wild-type ATRX, but not in a mutant lacking RNA binding ability^43^. Thus, it is plausible that TERRA recruits chromatin-modulating factors via direct binding with its partner proteins, implicating TERRA in 3D chromatin organization. Furthermore, 3D telomere looping^44-46^ may bring telomeres much closer to TERRA target loci, allowing TERRA to travel only short distances and facilitating its reach to other areas of the genome that are millions of base pairs away. This is an active area of research, and future studies may shed more light on the effect of telomere shortening on 3D chromatin organization.

Interestingly, while many lncRNAs have been shown to act via cis/trans binding to their DNA targets^47^, only one study to date has reported binding of a lncRNA in an R-loop associated manner to mediate the expression of target genes in trans^14^. This lncRNA, APOLO, was shown in Arabidopsis to decoy and displace protein complexes from its target sites and form R-loops via complementary base-pairing, allowing modulation of the 3D chromatin conformation, which affected gene transcription^14^. Another study showed that TERRA can bind telomeres via R-loops^13^. In this study by Feretzaki et al., cells were transiently transfected with plasmids encoding the sub-telomeric regions from selected chromosomes followed by 90 TTAGGG repeats, and showed that upon expression, these recombinant chimeric TERRA bind telomeric DNA in a TTAGGG-dependent manner. This was also confirmed using CRISPR-Cas9 integration of the chimeric constructs to achieve physiological expression levels^13^. This constitutes evidence that a recombinant chimeric TERRA can bind telomeres from which it did not originate, lending support to our findings that focus on the binding mechanism of TERRA to non-telomeric regions.

Additionally, we show that the RNA helicase ATRX is present at sites where TERRA is found. This aligns with previous work showing that TERRA and ATRX share many target genes, and that they are functionally antagonistic, as ATRX competes with TERRA for DNA binding^7^. The presence of ATRX at TERRA sites supports our conclusion that the R-loops present within a subset of TERRA peaks are the result of the presence of TERRA rather than being co-transcriptional or due to other non-TERRA related processes.

It is interesting that the majority of TERRA peaks lacked tandem repeats, which may hint at diverse mechanisms by which TERRA can perform its functional roles. Future studies should further explore non-telomeric TERRA functions that do not depend on the presence of tandem telomeric repeats.

Of further interest was our finding that the majority of R-loop-intersecting TERRA peaks (∼62%) do not contain telomeric repeats (Fig. 2B). We therefore cannot exclude the likelihood of TERRA RNAs forming R-loops at genomic regions that lack telomeric repeats. One explanation may be the presence of AG/TC-rich repeats within these regions (Fig. 2H and S2), which may partially mimic the G-rich telomeric repeat, providing a weak template for complementary binding and R-loop formation. Another explanation might be that these genomic regions share high sequence similarity with the sub-telomeric portions of many TERRA RNAs, enabling them to more easily form stable R-loops via complementary base-pairing. Furthermore, AG/TC-rich regions may support R-loop formation independent of TERRA, as was the case in several such genomic regions (Fig. S2), especially since R-loop presence was not exclusive to telomeric repeat-containing peak regions (Fig. S3A). TERRA presence at those regions may be due to association with other TERRA-binding proteins (e.g., adapter proteins or transcription factors) that in turn bind these DNA regions, meaning that TERRA may not directly be associated with DNA at these repeat-lacking peaks. This is supported by our observation of a 10-fold higher TERRA enrichment in tandem repeat-containing peak regions (Fig. 2G and 3B). Consequently, a portion of these R-loop peaks may be TERRA-independent.

When assessing intersecting TERRA peaks for TERRA-dependent gene expression, we expected a higher number of genes to overlap between both groups. However, these results may have been affected by several factors, both experimental and biological. One such factor may have been the difficulty of completely knocking down TERRA (and the infeasibility of knocking it out due to the essential nature of telomeres), as our knowledge is incomplete on whether a small amount of TERRA can still exert a significant effect. It could also be that the effect of TERRA knockdown requires a longer time period to be fully detected, and the experimental setup may not have allowed for this. Additionally, the authors^7^ only reported genes with a fold change greater than two, suggesting there may be a higher overlap with our TERRA-associated genes, albeit with a lower effect on gene expression. Furthermore, it is not yet known whether the binding dynamics of short and long TERRA molecules are similar, and how this would affect gene transcription and other TERRA-related functions as the cell progresses through many replication cycles, especially since telomeres on some chromosomes are shorter than others. It could be that TERRA’s effect on target gene expression increases with age as telomeres shorten. Future experimental setups should consider the age of the model cell/organism and the cellular state of telomeres, the fraction of short/long TERRA molecules, and whether sub-telomeric TERRA regions add more specificity to TERRA binding sites.

In conclusion, our work suggests a model by which TERRA acts as a telomeric second messenger, modulating gene transcription in trans via R-loops upon its transcription at telomeres, thereby extending the genomic reach of telomeres. This opens new avenues of research into how telomeres can modulate transcriptional landscapes spatially and temporally as cells age. Moreover, our findings raise the possibility that similar R-loop–mediated mechanisms may be broadly employed by other trans-acting lncRNAs to regulate the expression of their target genes.

## DATA AVAILABILITY

All scripts and code used in this study have been deposited in the following GitHub repository: https://github.com/saifshehata/terra_rloop/tree/master

## DECLARATION OF GENERATIVE AI AND AI-ASSISTED TECHNOLOGIES IN THE WRITING PROCESS

During the preparation of this work, the authors used ChatGPT (OpenAI) in order to refine scientific writing, improve clarity, and ensure consistency in the language of the manuscript. After using this tool/service, the author(s) reviewed and edited the content as needed and take(s) full responsibility for the content of the publication.

## ACKNOWLEDGEMENT

The computations and data handling were enabled by resources in project SNIC-2022-22-85 provided by the Swedish National Infrastructure for Computing (SNIC) at UPPMAX, partially funded by the Swedish Council through grant agreement no. 2018-05973. We acknowledge Anders Sjölander for assistance concerning technical and implementational aspects of the UPPMAX resources.

## FIGURE LEGENDS

**Supplementary figure**

**1)** Permutation test using randomized peak regions to assess the significance of overlap between TERRA and R-loop peaks.

**2)** Genome browser views of TERRA, R-loop, and ATRX coverage at intronic peaks containing fewer than four tandem telomeric repeats.

**3A)** Metagene profiles and heatmaps of R-loop coverage at intersecting peaks located in intronic or intergenic regions. Peaks were divided into groups with fewer than four versus at least four tandem telomeric repeats.

**3B–C)** Genome browser views of TERRA, R-loop, and ATRX coverage flanked by broad R-loop peaks in intronic regions with tandem telomeric repeats.

