## Supplementary figures and images for "Telomeric Repeat-Containing lncRNA TERRA Targets Non-Telomeric DNA in Trans via R-Loops"

### Supplementary figures

Fig. S1

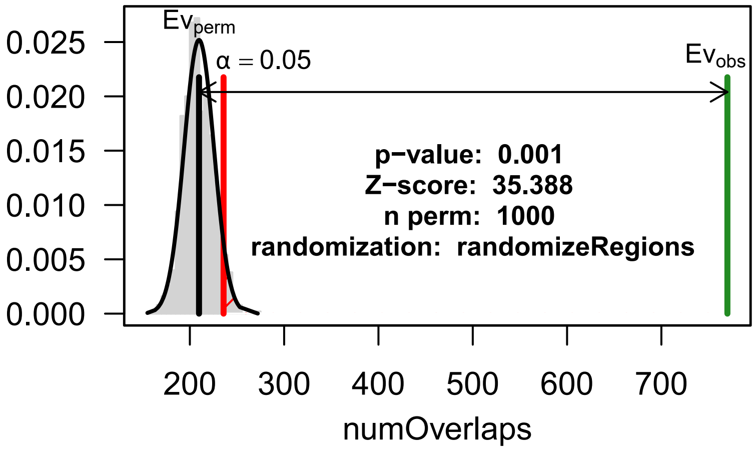


Fig. S2

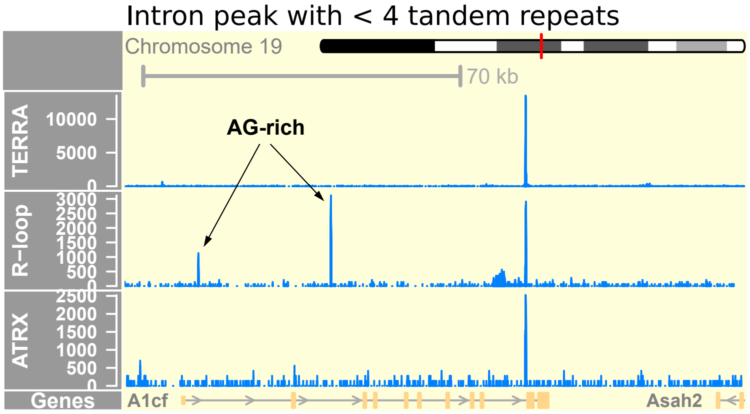


Fig. S3A

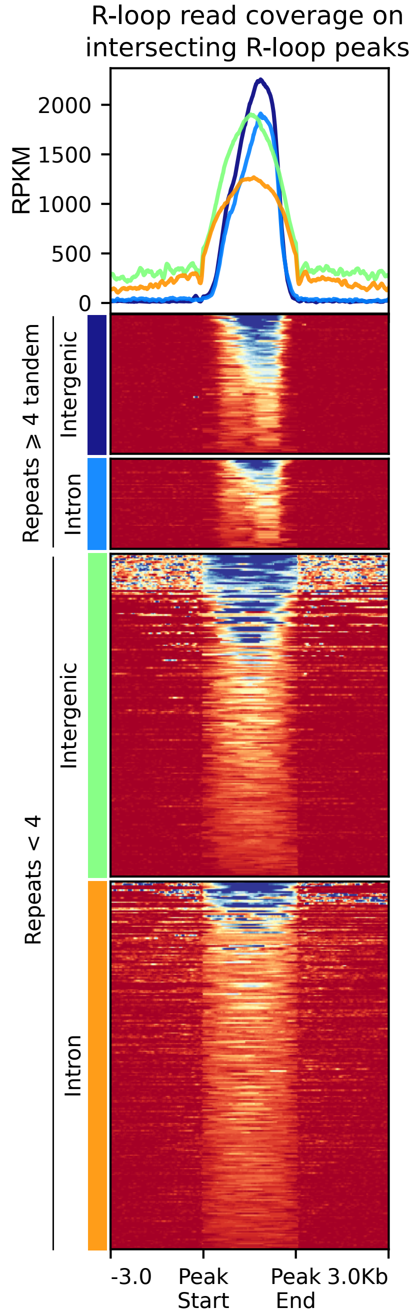


Fig. S3B

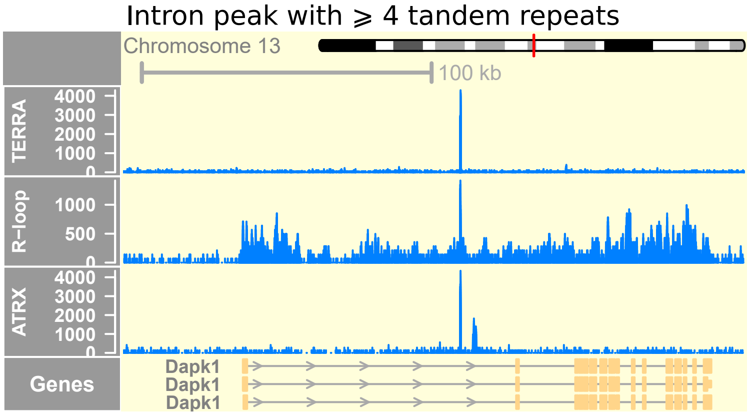


Fig. S3C

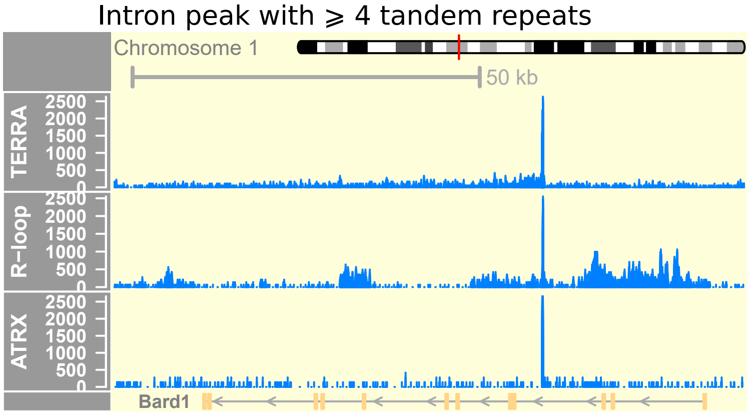
